## Supplementary data for "Prediction of A. thaliana’s MCTP4 Structure using Deep Learning-Based tools and Exploration of Transmembrane domain Dynamics using Coarse-Grained Molecular Dynamics Simulations"

Table 1: Statistical information on each model from OPM server.[1]

| Models | Tilt | Depth |
|---|---|---|
| AF | 15 ± 0° | 27.2 ± 1.2 Å |
| AFM | 40 ± 0° | 31.2 ± 0.7 Å |
| OF | 37 ± 4° | 31.2 ± 0.8 Å |
| ESM | 34 ± 0° | 27.8 ± 2.4 Å |
| RF | 72 ± 1° | 17.8 ± 1.2 Å |
| TR | 33 ± 2° | 30.8 ± 1.1 Å |

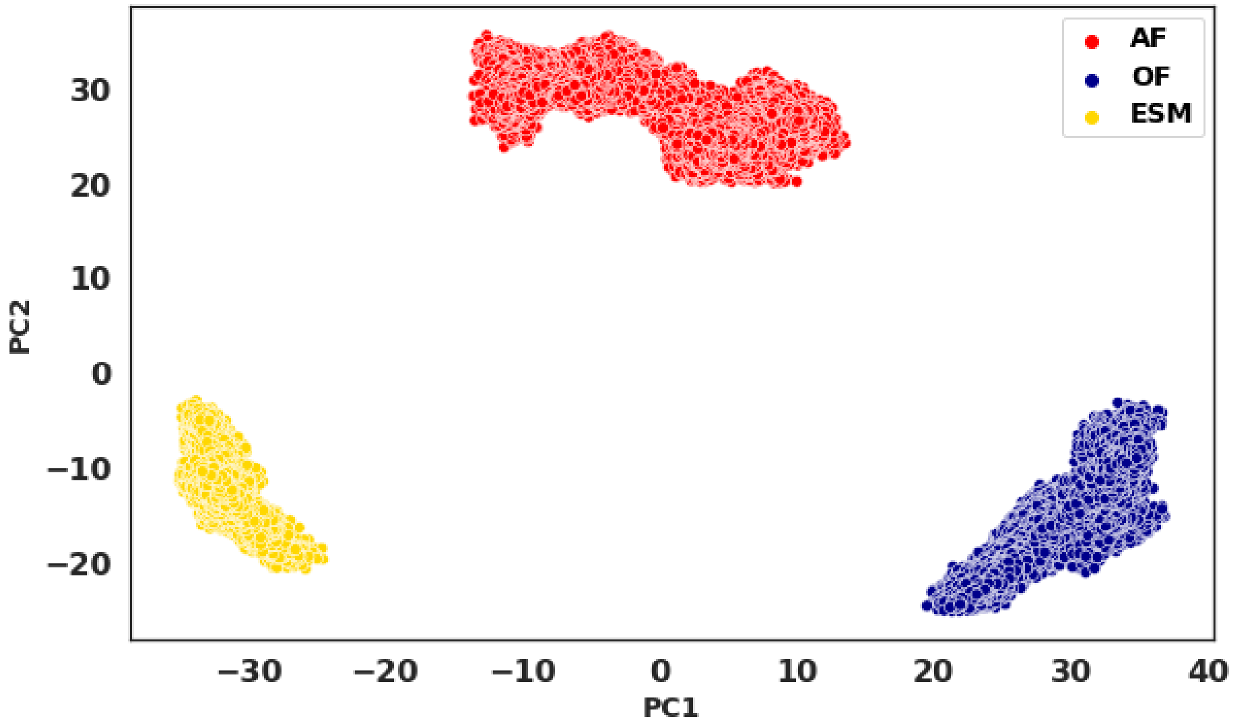


(a)

Figure 1: Projection of the first principal components (PC1 against PC2) from all atom simulation. Each point represents an observation. Colors represent different models: Red for AF2, Navy Blue for OMEGA, and Gold for ESM

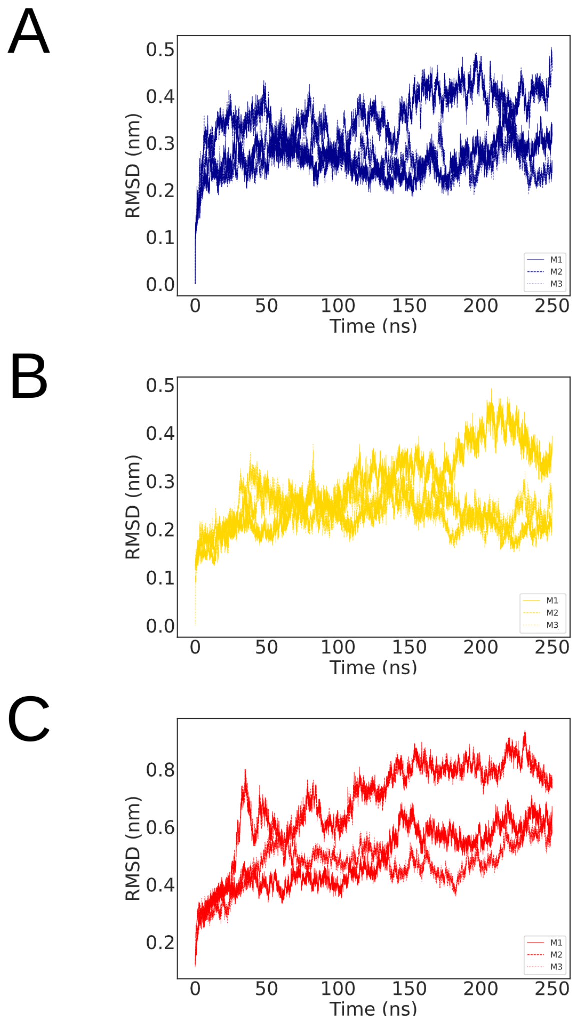


(a)

Figure 2: RSMD from all-atom simulation. Red for AF2, Navy Blue for OMEGA, and Gold for ESM

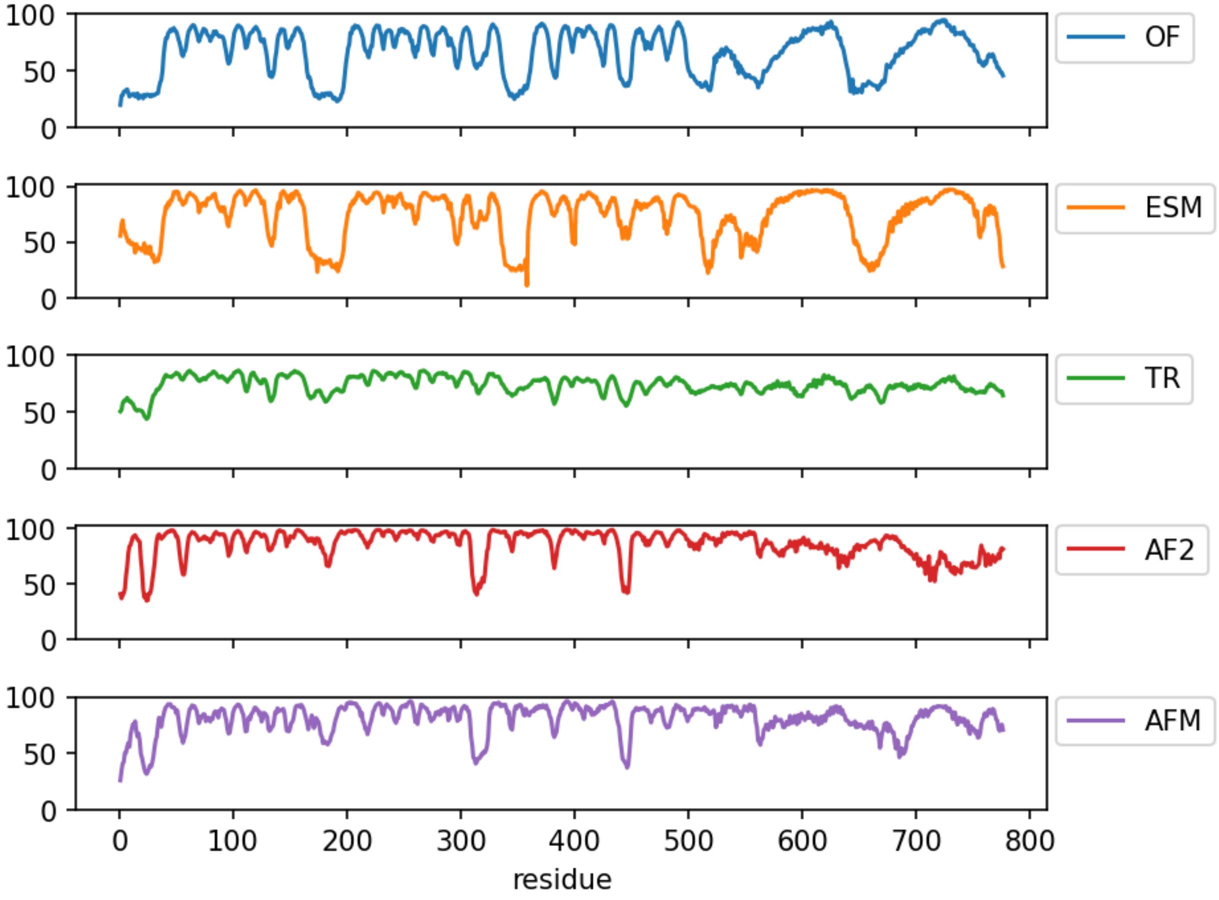


(a)

Figure 3: Evaluation of model predictions using the pLDDT score. The curves of different colors represent models predicted by various prediction methods: AlphaFold (AF, red), OmegaFold (OF, blue), TR (green), ESM (orange), and AlphaFold Multimer (AFM, purple)

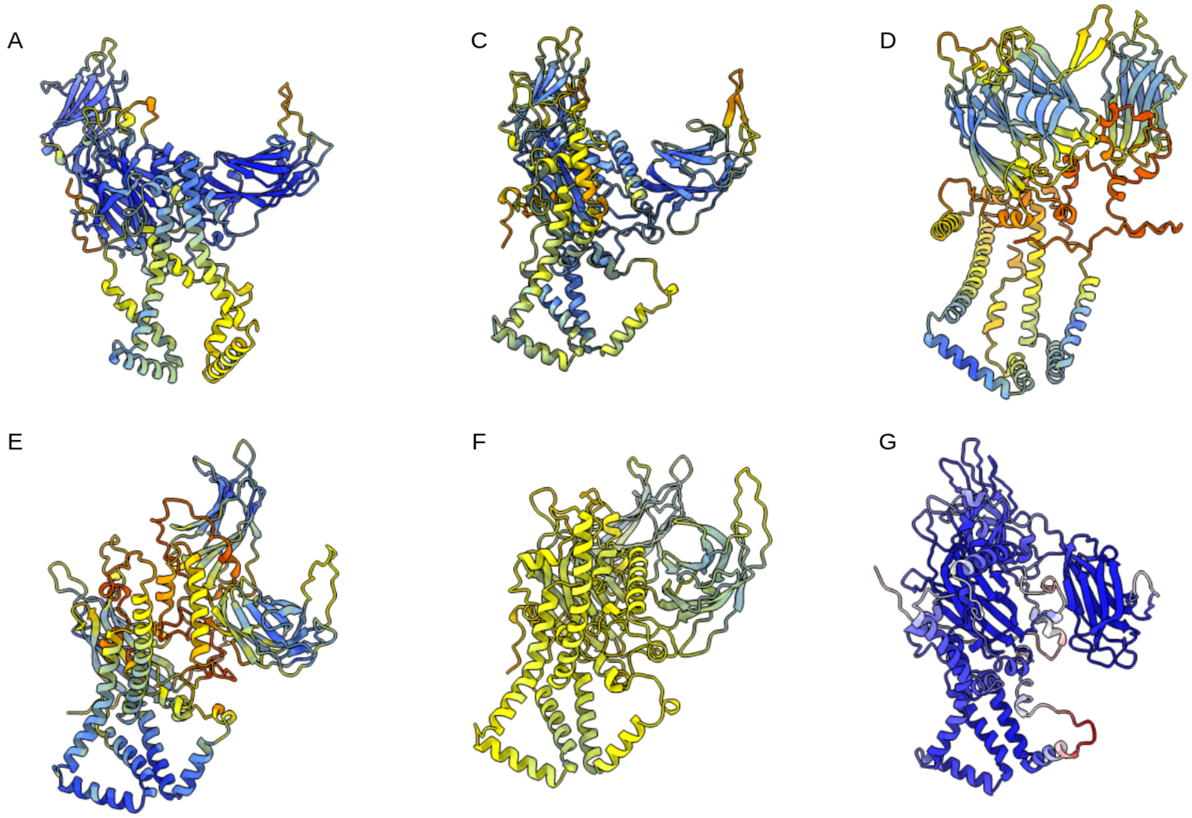


(a)

Figure 4: Panels A to F (AF, AFM, OF, ESM, TR): Structures are colored according to pLDDT scores; blue signifies areas with high confidence (reliable structural prediction), while red depicts areas with low confidence (uncertain structural prediction). Panel G displays the prediction made by RosettaFold, colored on the error estimate in the Å (RMSD) metric. This metric categorizes structural predictions into three levels of confidence: blue for very confident, red for less confident, and white for areas lacking confidence.

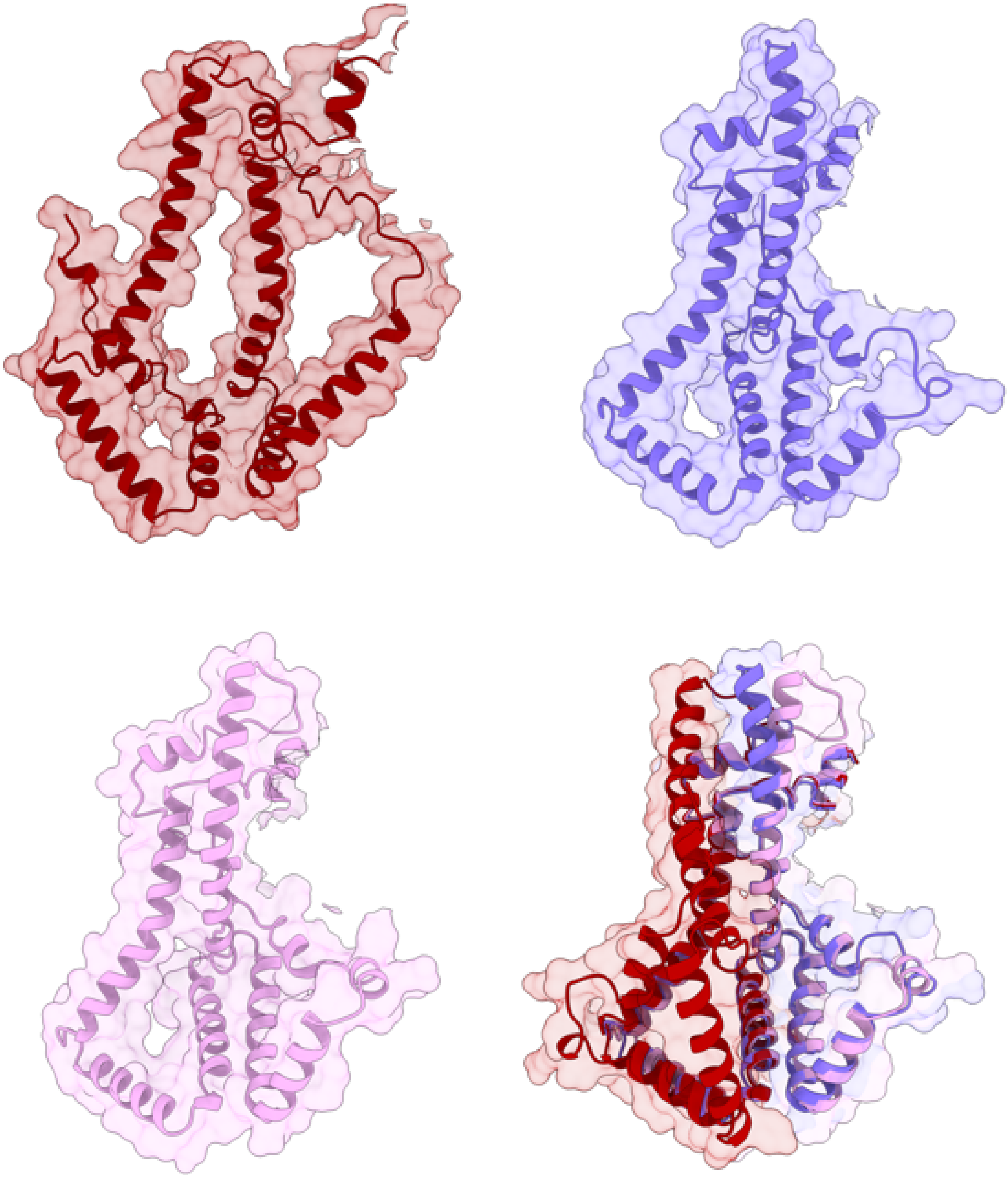

(a)

Figure 5: Models from AF3 (dark red), Boltz-1 (light purple) and Chai-1 (light pink) plotted on the PCA landscape. The AF3 model shows structural TMR similarities with the Omegafold model, while Boltz-1 and Chai-1 align with the consensus conformations observed in ESM, AFM, TR, and RF models.

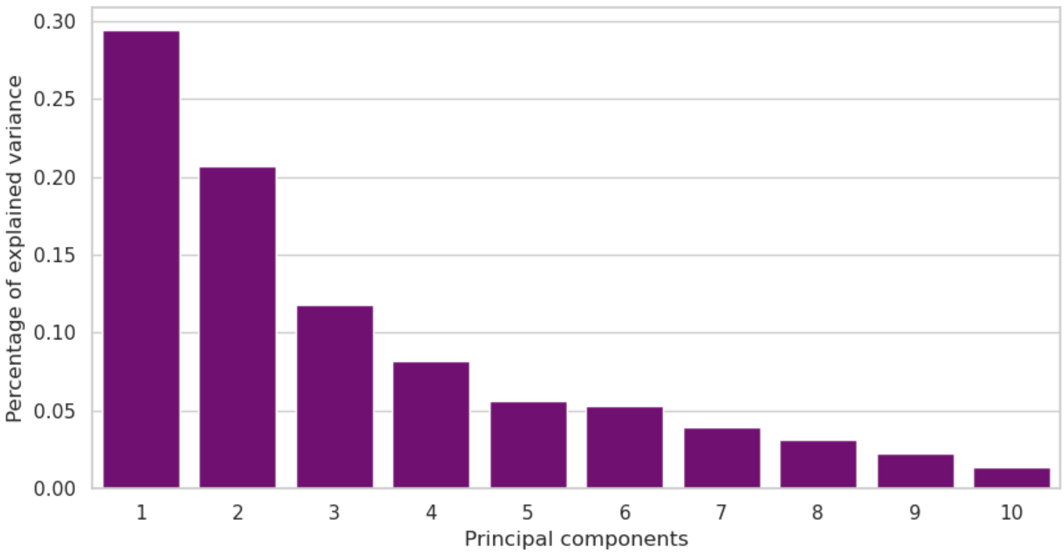


(a)

Figure 6: Variance explained by each Principal Component (PC) on CG simulations).

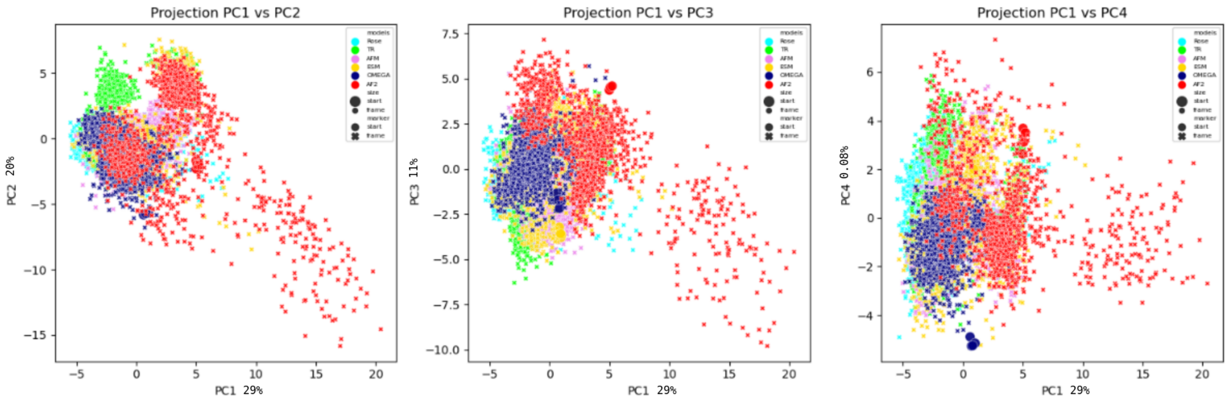


(a)

Figure 7: Projection of the first principal components (PC1 against PC2, PC3 and PC4). Each point represents an observation. Colors represent different models: Aqua for Rose, Lime Green for TR, Violet for AFM, Gold for ESM, Navy Blue for OMEGA, and Red for AF2

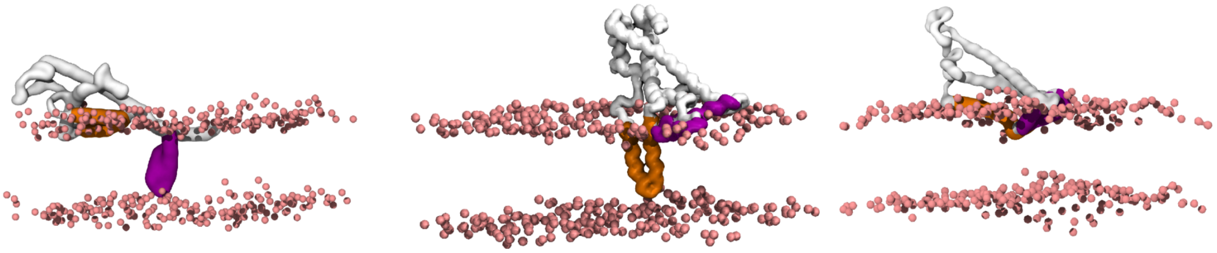

(a)

Figure 8: Left: HP2 domain emerging from the membrane. Center: HP1 domain emerging from the membrane. Right: Both HP1 and HP2 domains emerging from the membrane. These emergences from the membrane are observed in some replicas, regardless of the system in CG simulations. For our analysis, we chose not to consider these instances. Other predictive tools, such as PSIPRED[2] or DREAMM,[3] indicate that this domain remains inside the membrane. When one or both domains emerge, they never re-enter the membrane during the simulation.

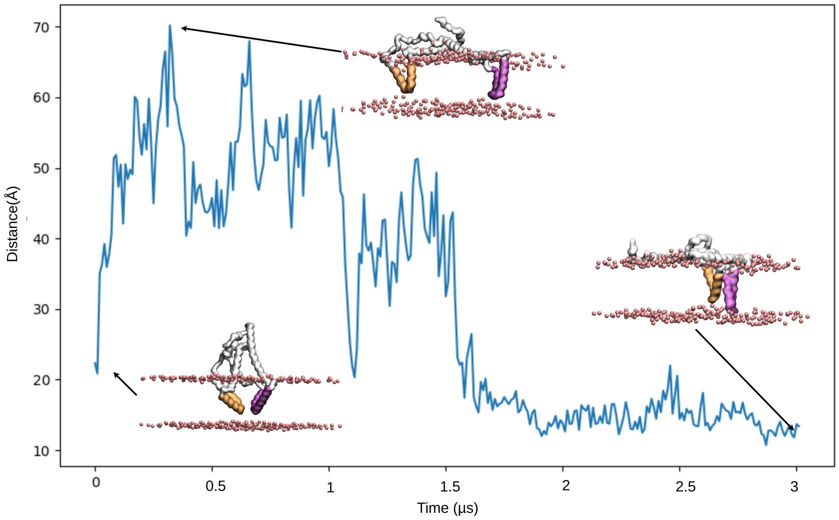


(a)

Figure 9: Distance between HP1 and HP2 domains throughout an AlphaFold CG simulation.

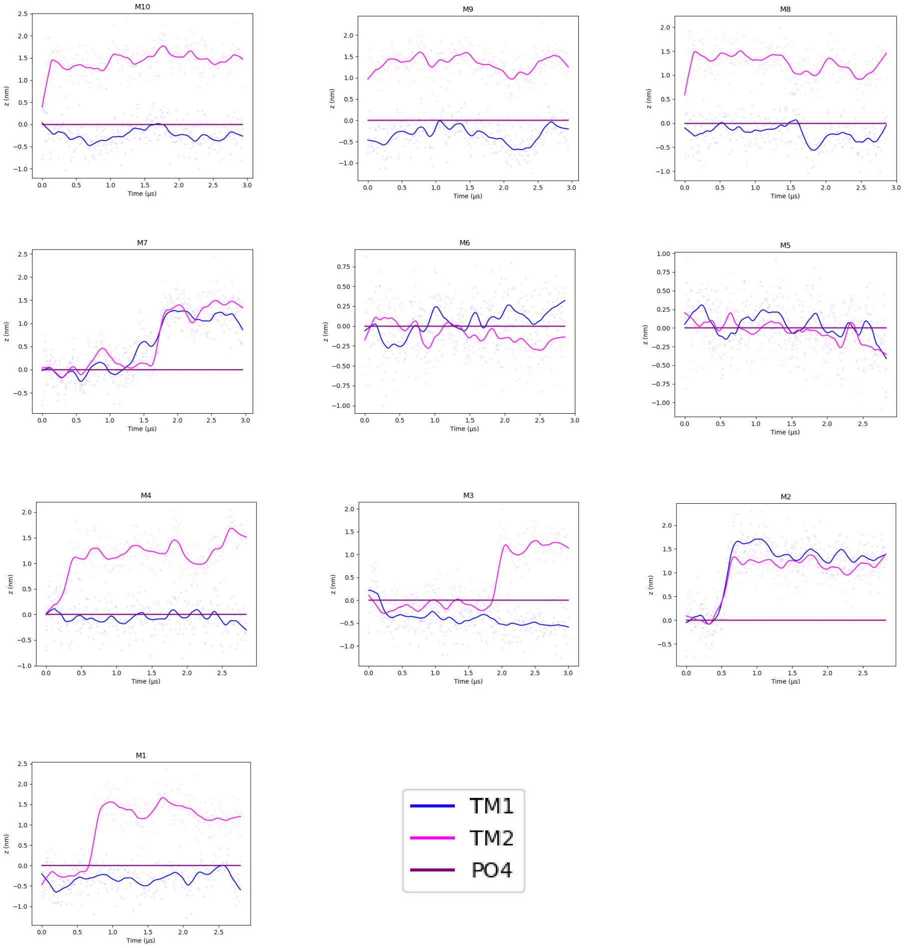


(a)

Figure 10: Distance of the center of mass along the Z-axis for the HP1 and HP2 domains of the AF model

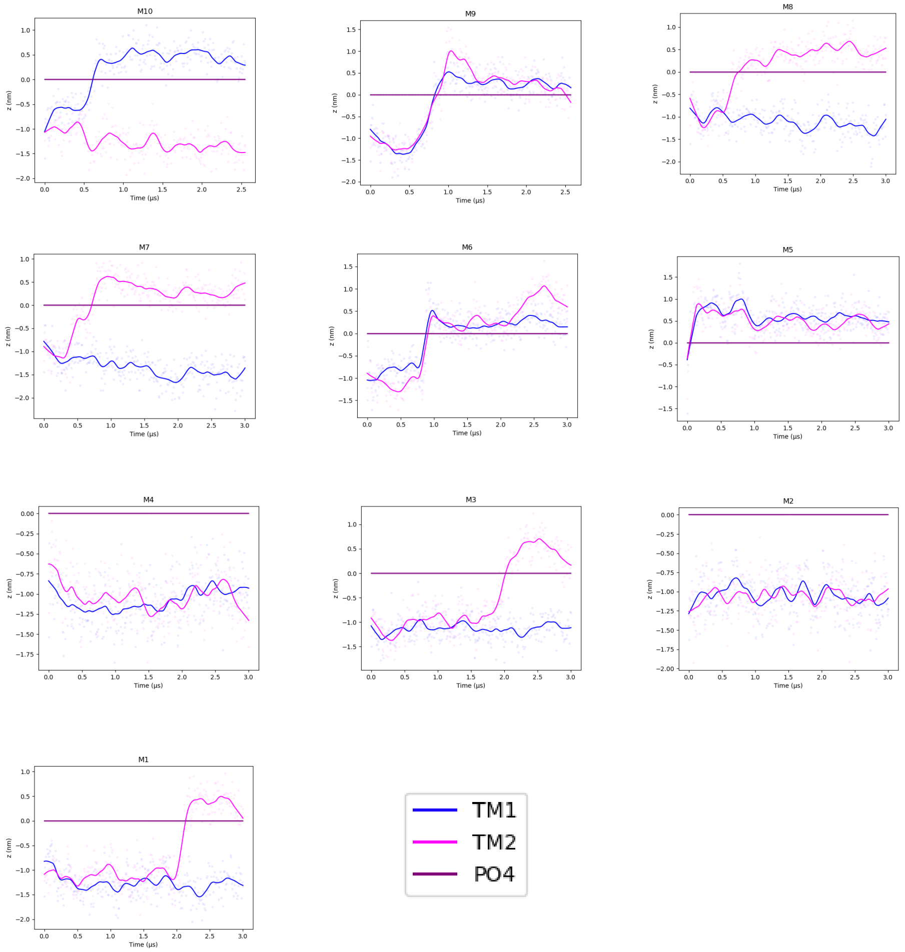


(a)

Figure 11: Distance of the center of mass along the Z-axis for the HP1 and HP2 domains of the AFM model

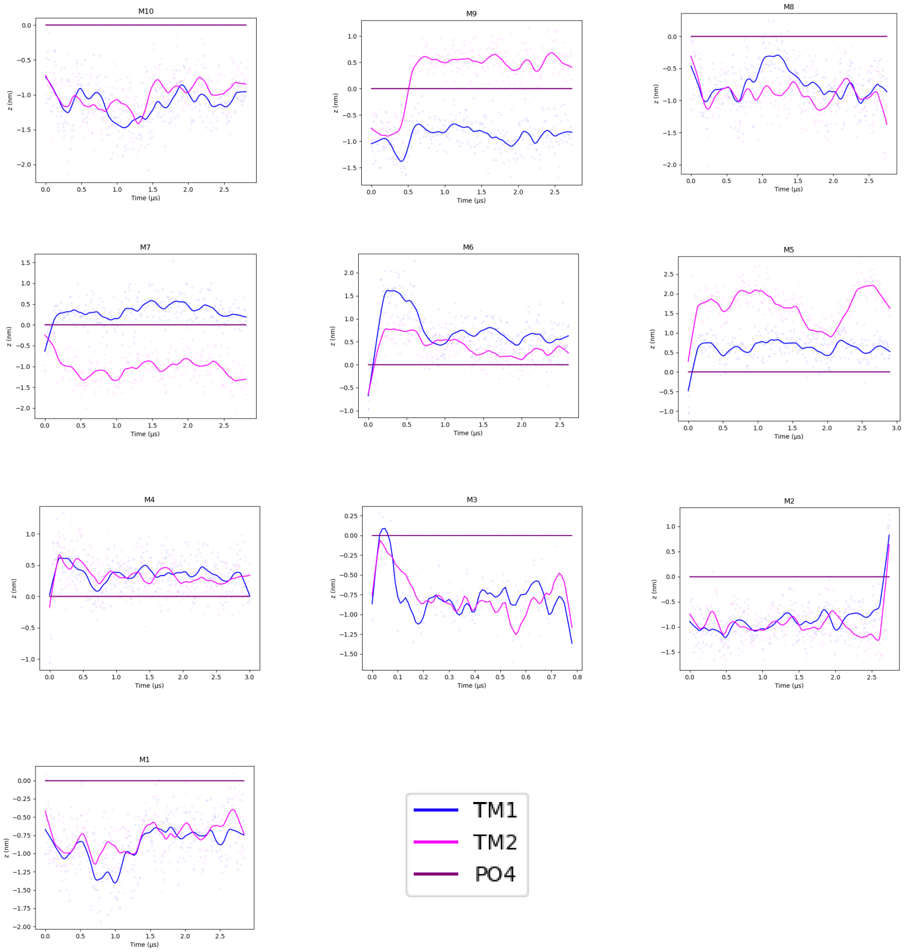


(a)

Figure 12: Distance of the center of mass along the Z-axis for the HP1 and HP2 domains of the RF model

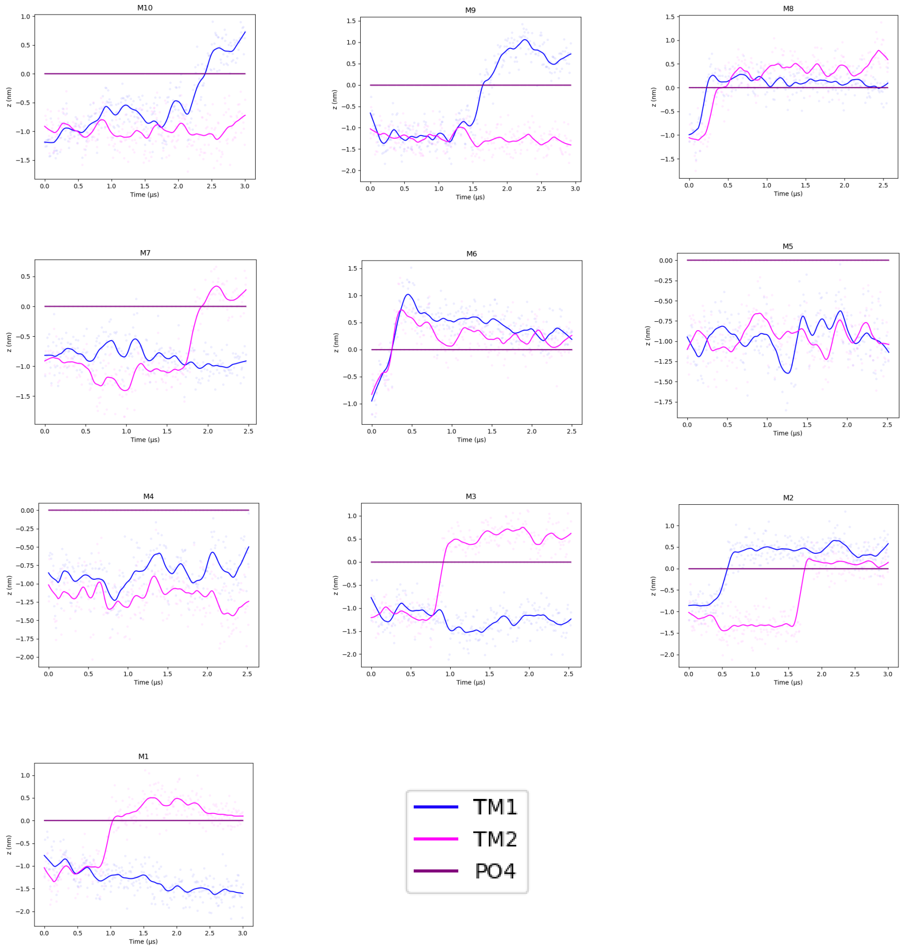


(a)

Figure 13: Distance of the center of mass along the Z-axis for the HP1 and HP2 domains of the TR model

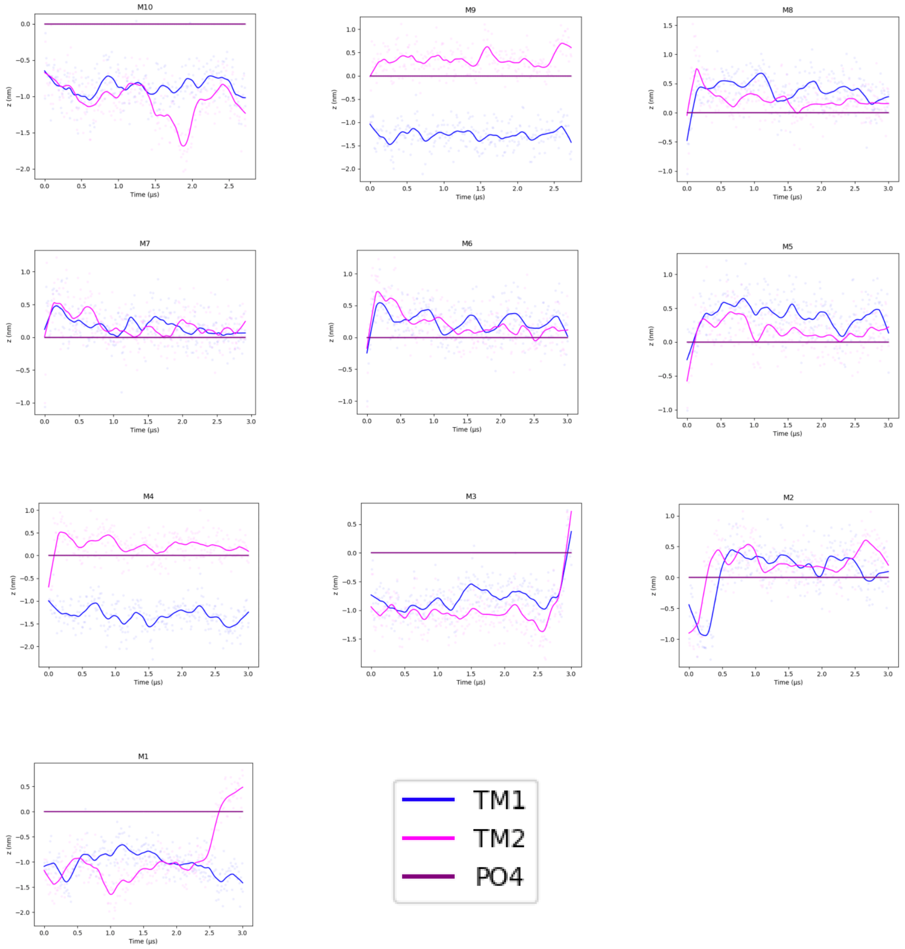


(a)

Figure 14: Distance of the center of mass along the Z-axis for the HP1 and HP2 domains of the OF model

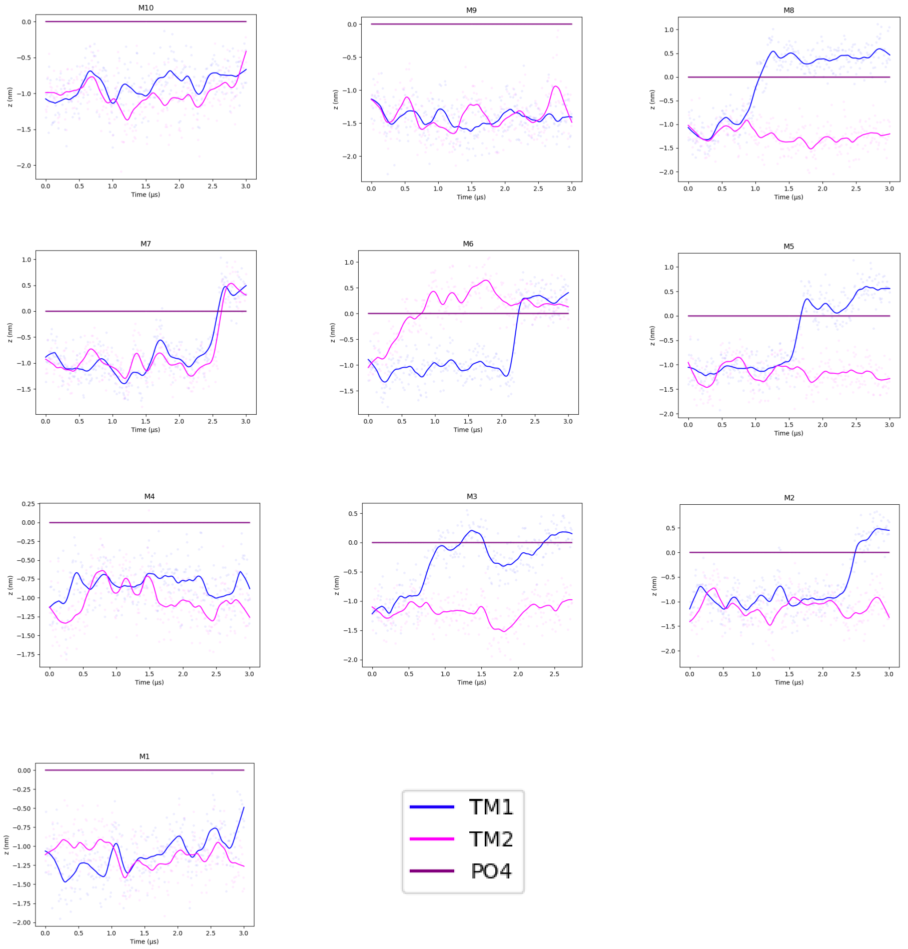


(a)

Figure 15: Distance of the center of mass along the Z-axis for the HP1 and HP2 domains of the ESM model

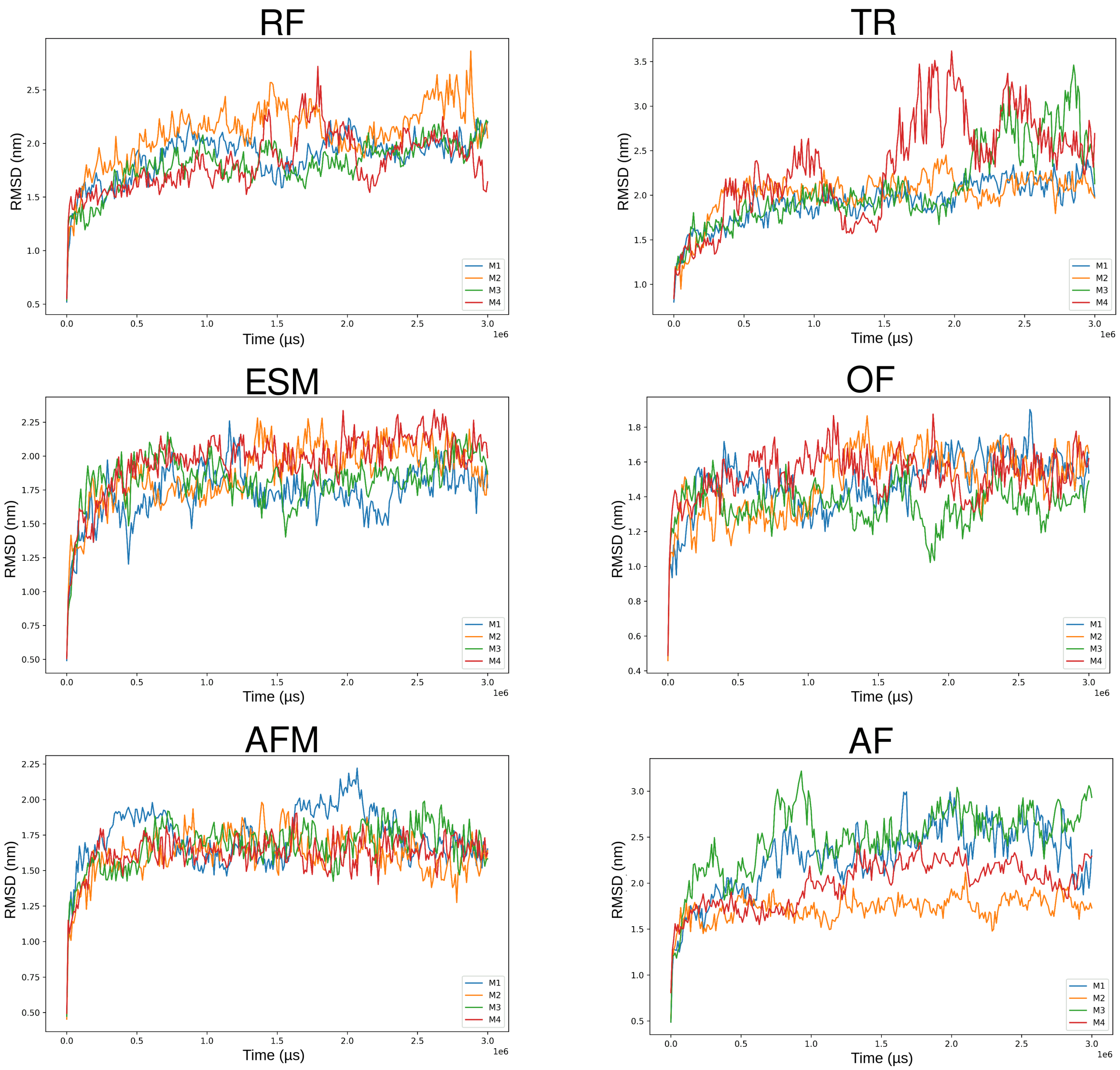


(a)

Figure 16: RMSD for CG simulations.
